# Pattern separation and tuning shift in human sensory cortex underlie fear memory

**DOI:** 10.1101/2021.08.27.457990

**Authors:** Yuqi You, Lucas R. Novak, Kevin Clancy, Wen Li

## Abstract

Animal research has recognized the role of the sensory cortex in fear memory and two key underlying mechanisms—pattern separation and tuning shift. We interrogated these mechanisms in the human sensory cortex in an olfactory differential conditioning study with a delayed (9-day) retention test. Combining affective appraisal and olfactory psychophysics with functional magnetic resonance imaging (fMRI) multivoxel pattern analysis and voxel-based tuning analysis over a linear odor-morphing continuum, we confirmed affective and perceptual learning and memory and demonstrated associative plasticity in the human olfactory (piriform) cortex. Specifically, the piriform cortex exhibited immediate and lasting enhancement in pattern separation (between the conditioned stimuli/CS and neighboring non-CS) and late-onset yet lasting tuning shift towards the CS, especially in anxious individuals. These findings highlight an evolutionarily conserved sensory cortical system of fear memory, which can underpin sensory encoding of fear/threat and confer a sensory mechanism to the neuropathophysiology of anxiety.

## INTRODUCTION

Aversive conditioning generates reliable fear or threat learning and memory, providing prominent experimental models of anxiety and depression ^1, 2, 3, 4^. Beyond the well-established role of the amygdala and prefrontal cortex (PFC) in aversive conditioning, rapidly accruing evidence has expanded the “fear circuit” to include the sensory cortex ^3, 5, 6, 7, 8^. In this fear circuit, the amygdala supports fear acquisition and consolidation, the prelimbic cortex (or the human homologue—anterior cingulate cortex) and insula underpin fear orientation and attention, and the orbitofrontal/ventromedial PFC subserves fear extinction ^3, 9, 10^. As for the sensory cortex, recent discoveries and theories have underscored a role in the long-term fear memory ^7, 11, 12^.

As early as the 1950s ^13^, researchers have observed sensory cortical plasticity (e.g., enhanced response in the primary auditory cortex to the conditioned stimulus/CS) following aversive conditioning. Such *associative* sensory cortical plasticity not only emerges immediately after conditioning but also lasts for days to weeks ^14, 15^. Recent rodent evidence further indicates that associative sensory cortical plasticity plays a critical role in the formation ^16, 17, 18^ and storage of long-term memory of aversive conditioning ^11, 16, 19, 20, 21^. In humans, an increasing number of studies, from our group and others, have confirmed associative plasticity in the human sensory cortex following aversive conditioning ^22, 23, 24, 25, 26^. In support of long-term storage of aversive conditioning in the human sensory cortex, our group further demonstrated enhanced CS response in the human primary visual cortex (V1/V2) 15 days after conditioning ^27^. However, the mechanism underlying fear memory in the human sensory cortex remains unexplored.

Animal research has indicated that two key mnemonic mechanisms—pattern separation (supporting discrimination among similar cues) and pattern completion (permitting memory activation by partial cues)—can support long-term memory of conditioning in the sensory (particularly, olfactory) cortex ^28, 29^. The primary olfactory cortex (i.e., piriform cortex) is thought to resemble an associative, content-addressable memory system, ideally positioned to subserve long-term memory of conditioning ^8, 30, 31^. Indeed, olfactory associative plasticity, including pattern separation (between CS and similar odors) and completion (between CS and dissimilar cues), has been observed across phylogeny ^8, 32, 33, 34^. In humans, olfactory differential conditioning has been found to induce immediate pattern separation in the piriform cortex ^22^ and enhance perceptual discrimination between the CS and similar odors ^22, 35, 36, 37^. Therefore, we hypothesize that this pattern separation in the human sensory cortex could persist to underpin long-term fear memory.

Animal research has further implicated “associative representational plasticity” as a mechanism underlying long-term memory and sharpened perception of the CS ^12^. This associative representational plasticity is characterized by tuning shift in the sensory cortex such that neurons initially tuned to non-CS shift their tuning to respond maximally to the CS ^14, 15, 33, 38^. Importantly, such associative representational plasticity/tuning shift would consolidate over time and last for a long time, thereby underpinning stable sensory representation and long-term memory of the CS ^6, 12, 38, 39^. As indirect evidence of associative tuning shift in the human sensory cortex, a human electrophysiological study demonstrated that visual cortical responses were both enhanced for the CS and suppressed for the most similar non-CS ^26^. It is thus plausible that associative tuning shift can also occur and persist in the human sensory cortex to support long-term fear memory. We thus examined pattern separation and tuning shift in the human olfactory cortex (anterior and posterior piriform cortices, APC/PPC) using olfactory aversive conditioning with delayed (9-day) retention tests. As in previous animal ^12, 40, 41^ and human ^26^ research, to elucidate these mechanisms, we included a linear morphing continuum of odor mixtures in an odor discrimination task (ODT), with the two extreme odor mixtures differentially paired with aversive and neutral unconditioned stimuli (i.e., threat CS/CSt and safety CS/CSs, respectively; Fig. 1a-c). Pattern separation and tuning shift in the primary olfactory cortex was assessed with fMRI multivoxel representational similarity analysis (RSA) ^22^ and fMRI voxel-based tuning analysis ^42, 43^, respectively. For comparison, supplemental analyses of these processes were performed in the amygdala and orbitofrontal cortex (OFC), key substrates of the classical fear circuit in (Fig. 1d). Importantly, akin to fear conditioning models of anxiety, aversive conditioning is amplified by anxiety ^44, 45^, and particularly relevant here, sensory perceptual effects of aversive conditioning could be heightened by anxiety ^27, 46^. We thus examined the modulatory effects of anxiety on these mechanisms.

**Fig. 1.**
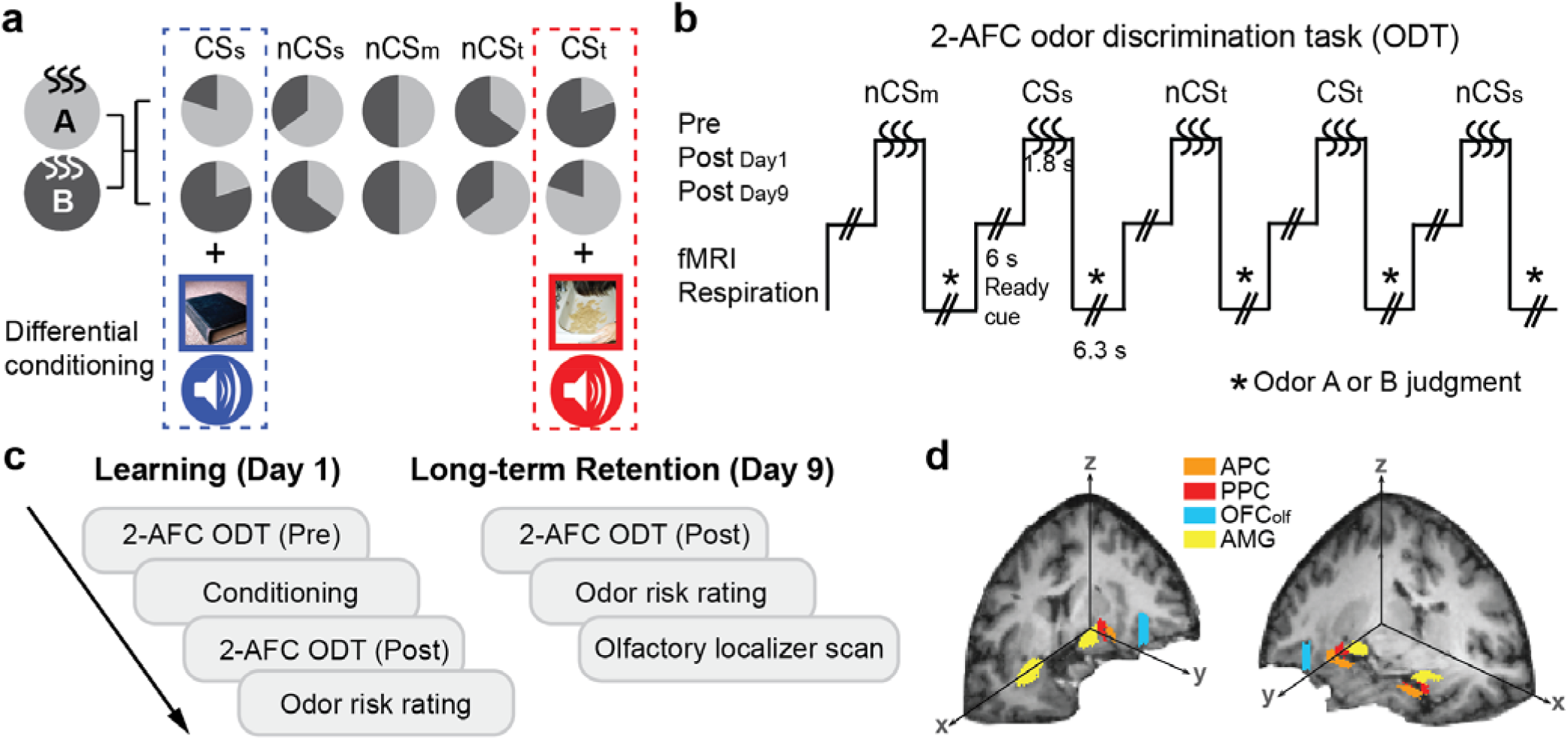
Odor stimuli and experimental design. (a) Stimuli consisted of a continuum of five parametrically-morphed binary odor mixtures of neutral odors (acetophenone and eugenol labeled as Odor A and Odor B). The extreme mixtures (20% A/80% B and 80% A/20% B) were differentially conditioned as CSt (threat) or CSs (safety) via paired presentation with aversive or neutral unconditioned stimuli (UCS: bimodal aversive or neutral pictures and sounds). Assignment of CSt/CSs was counterbalanced across participants. The three intermediate mixtures (35% A/65% B, 50% A/50% B, and 65%A /35% B) were non-conditioned stimuli (nCS), representing the odor neighboring CSt (nCSt), the midpoint mixture (nCSm), and the odor neighboring CSs (nCSs). (b) Two-alternative-forced-choice (2-AFC) odor discrimination task (ODT) accompanied by fMRI and respiration acquisition. Each trial presented an odor mixture pseudo-randomly for 1.8 seconds, to which participants made judgments of “Odor A” or “Odor B” with button pressing. (c) Experiment schedule. Day 1 consisted of pre-conditioning 2-AFC ODT, conditioning, post-conditioning 2-AFC ODT, and odor risk rating. Day 9 consisted of post-conditioning 2-AFC ODT, odor risk rating, and an olfactory localizer scan. (d) Regions of interest (ROIs). Anatomical masks of the primary olfactory cortex (anterior piriform cortex/APC and posterior piriform cortex/PPC), the olfactory orbitofrontal cortex (OFC_olf_), and the amygdala (AMG) are displayed on 3D T1 sections of one participant. These ROIs were further functionally constrained by the olfactory localizer.

## MATERIALS AND METHODS

### Participants

Thirty-three individuals (13 males; age 19.9 ± 2.0 years, range 18–25) participated in this two-session fMRI experiment in exchange for course credit or monetary compensation. All participants were right-handed, with normal olfaction and normal or corrected-to-normal vision. Participants were screened to exclude acute nasal infections or allergies affecting olfaction, any history of severe head injury, psychological/neurological disorders, or current use of psychotropic medication. All participants provided informed consent to participate in the study, which was approved by the University of Wisconsin-Madison Institutional Review Board. One participant who failed to provide risk ratings on Day 1 and another who failed to follow the ODT task instruction were excluded from the corresponding analyses. Two participants were excluded from fMRI analysis due to metal artefact and excessive movement.

### Anxiety Assessment

We used the Behavioral Inhibition Scale (BIS) to measure trait anxiety^47^. The BIS is a 7-item self-report questionnaire (score range: 7-28) measuring the strength of the behavioral inhibition system and threat sensitivity, known to reflect trait anxiety. This scale is neurobiologically motivated, with high reliability and strong predictive validity of anxiety ^45, 48^, and recommended by the National Institute of Mental Health (NIMH) to measure the construct of “potential threat (anxiety)”.

### Stimuli

We included two neutral odorants, acetophenone (5% l/l; diluted in mineral oil) and eugenol (18% l/l). These odors have received similar ratings on valence, intensity, familiarity, and pungency and been used as neutral odors in previous research ^49, 50^. They were labeled as odors “A” and “B” to the participants and were parametrically mixed into five mixtures to create a linear morphing continuum: 80% A/20% B, 65% A/35% B, 50% A/50% B, 35% A/65% B, and 20% A/80% B (Fig. 1a). The two extreme mixtures (20% A/80% B and 80% A/20% B) served as conditioned stimuli (CS), differentially conditioned as CS-threat (CSt) and CS-safety (CSs), counterbalanced across participants, via pairings with threat and neutral unconditioned stimuli (UCS), respectively. The three intermediate mixtures were non-conditioned stimuli (nCS) and denoted as nCSt (neighboring odor of the CSt), nCSm (midpoint of the continuum), and nCSs (neighboring odor of the CSs), respectively. The UCS were bimodal (visuo-auditory) stimuli, including 7 pairs of disgust images (three depicting dirty toilets and four vomits) and disgust sounds (i.e., vomiting) and 7 pairs of neutral images (household objects) and neutral sounds. Images were chosen from the International Affective Picture Set (IAPS) ^51^ and internet sources ^52^. Disgust sounds were from the disgust subset of human affective vocalizations ^53^, and neutral sounds were pure tones (300, 500, and 800 Hz).

Odor stimuli were delivered at room temperature using an MRI-compatible sixteen-channel computer-controlled olfactometer (airflow set at 1.5 L/min), which permits rapid odor delivery in the absence of tactile, thermal or auditory confounds ^49, 54, 55, 56^. Stimulus presentation and collection of responses were controlled using Cogent2000 software (Wellcome Department of Imaging Neuroscience, London, UK) as implemented in Matlab (Mathworks, Natick, MA).

### Two-alternative forced-choice odor discrimination task (2-AFC ODT)

During the 2-AFC ODT, each trial began with a visual “Get Ready” cue, followed by a 3-2-1 countdown and a sniffing cue, upon which participants were to take a steady and consistent sniff and respond whether the odor smelled like Odor A or B by button pressing (Fig. 1b). Each of the five odor mixtures was presented 15 times, in a pseudo-random order without repetition over two consecutive trials. Seven additional trials with a central, blank rectangle on the screen (no response required) were randomly intermixed with the odor trials to help minimize olfactory fatigue and establish a non-odor fMRI baseline. Trials recurred with a stimulus onset asynchrony of 14.1 s.

### Experiment procedure

#### Pre-experiment screening

Approximately a week before the experiment, participants visited the lab to be screened for normal olfactory perception. They were also introduced to acetophenone and eugenol as Odors “A” and “B” and practiced on a 2-AFC ODT between the two odors. They also provided ratings on the five odor mixtures (see Supplemental Results).

#### Experiment Day 1

Participants first performed the 2-AFC ODT, then underwent differential conditioning, and then repeated the 2-AFC ODT (Fig. 1c). During differential conditioning, CSt and CSs odors were presented (seven trials each, randomly intermixed) for 1.8 s while the aversive or neutral UCS were presented respectively for 1.5 s at 1 s after CS odor onset, with 100% contingency. To prevent extinction by the repeated unreinforced CS presentation during the post-conditioning 2-AFC ODT (on both Day 1 and Day 9), five extra trials of CSt paired with the aversive UCS were randomly inserted ^22, 23, 27, 57^. Data from these trials were excluded from analysis. After the post-conditioning ODT, the five odor mixtures were presented (three trials per odor mixture, randomly intermixed), to which participants performed risk rating (likelihood of an aversive UCS to follow the odor) on a visual analog scale (VAS; 0-100%).

#### Experiment Day 9

Participants repeated the 2-AFC ODT and risk rating. After that, participants underwent an independent olfactory localizer scan involving a simple odor detection task, from which functional ROIs were extracted. Four additional odorants (α-ionone, citronellol, methyl cedryl ketone, 2-methoxy-4-methylphenol), neutral in valence and matched for intensity, were presented (15 trials/odor), pseudo-randomly intermixed with 30 air-only trials.

### Respiratory monitoring

Respiration measurements were acquired (1000 Hz) during the ODT, using a BioPac MP150 with a breathing belt affixed to the participant’s chest to record abdominal or thoracic contraction and expansion. For each odor trial, a sniff waveform was extracted from a 6 s window post sniff onset and was baseline-corrected by subtracting the mean activity within 1 s preceding sniff onset. Sniff parameters (inspiratory volume, peak amplitude, and peak latency) were generated by averaging across all 15 trials per odor. No odor effects were observed in these sniff parameters (see Supplemental Results).

### Imaging acquisition and preprocessing

Gradient-echo T2 weighted echoplanar images (EPI) were acquired with blood-oxygen-level-dependent (BOLD) contrast and sagittal acquisition on a 3T GE MR750 MRI scanner. Imaging parameters were TR/TE = 2350/20 ms; flip angle = 60°, field of view = 220 mm, slice thickness = 2 mm, gap = 1 mm; in-plane resolution/voxel size = 1.72×1.72 mm; matrix size = 128×128. A field map was acquired with a gradient echo sequence, which was coregistered with EPI images to correct EPI distortions due to susceptibility. A high-resolution (1×1×1mm^3^) T1-weighted anatomical scan was acquired. Five scan runs, including pre-conditioning, conditioning, Day 1 post-conditioning, Day 9 post-conditioning, and odor localizer, were acquired. Six “dummy” scans from the beginning of each scan run were discarded in order to allow stabilization of longitudinal magnetization. Imaging data were preprocessed in SPM12 (www.fil.ion.ucl.ac.uk/spm), where EPI images were slice-time corrected, realigned, and field-map corrected. Images collected on both Day 1 and Day 9 sessions were spatially realigned to the first image of the first scan run on Day 1, while the high-resolution T1-weighted scan was co-registered to the averaged EPI of both scan sessions. All multivariate pattern analyses were conducted on EPI data that were neither normalized nor smoothed to preserve signal information at the level of individual voxels, scans, and participants.

A general linear model (GLM) was computed on pre-conditioning ODT, conditioning, Day 1 post-conditioning ODT, and Day 9 post-conditioning ODT scans. Applying the Least Squares All (LSA) algorithm, we set each odor trial as a separate regressor, convolved with a canonical hemodynamic response function ^58^. Six movement-related regressors (derived from spatial realignment) were included to regress out motion-related variance. For the odor localizer scan, we applied a GLM with odor and no odor conditions as regressors, convolved with a canonical hemodynamic response function and the temporal and dispersion derivatives, besides the six motion regressors of no interest. A high-pass filter (cut-off, 128 s) was applied to remove low-frequency drifts and an autoregressive model (AR1) was applied to account for temporal nonsphericity.

### ROI definition

All four ROIs (APC, PPC, OFC, and amygdala) were manually drawn on each participant’s T1 image in MRIcro ^59^ (Fig. 1d). The olfactory OFC (OFC_olf_) was defined by a meta-analysis ^60^ and a prior study ^61^, and the other ROIs were defined by a human brain atlas ^62^. Left and right hemisphere counterparts were merged into a single ROI. Functional constraints were applied to these anatomical ROIs based on the odor-no-odor contrast of the independent odor localizer scan for each participant, with a liberal threshold at *P* < 0.5 uncorrected ^22^.

### fMRI analysis

#### Representational similarity analysis (RSA)

The RSA uses correlations across multivoxel response patterns to indicate the degree of similarity in response patterns ^63, 64^ and thus presents an effective test of pattern separation ^22^. For each participant and every ODT session, trial-wise beta values were extracted for all voxels within a functionally constrained ROI, which were then averaged across all 15 trials for each odor mixture, resulting in an odor-specific linear vector of beta values across a given ROI. Pearson’s correlation (*r*) was computed between all pairs of pattern vectors at each session, resulting in a 5 × 5 correlation matrix—the representational similarity matrix—for each session. To directly represent pattern separation, this matrix was converted into a representational dissimilarity matrix (RDM) by replacing the *r* values with dissimilarity scores (1 – *r*) ^65^. To assess pattern separation, we computed a pattern separation index (PSI) based on the RDM matrix (dissimilarity/distance = 1- *r*), following Fisher’s Z transformation: PSI = [(d1+ d4) – (d2 + d3)], reflecting the dissimilarity/distance of CSt and CSs from their neighboring nCS odors (nCSt and nCSs, d1 and d4 respectively), controlled by the dissimilarity/distance between the midpoint odor (nCSm) and its neighbors (d2 and d3).

#### Tuning analysis

We adopted a voxel-based tuning analysis used for visual sensory encoding ^42, 43^ to assess olfactory cortical tuning. Trial-wise beta values (5 odors×15 trials) for each voxel were normalized (by z-scoring) across trials after removing the trial-wise mean beta across the ROI, from which we calculated mutual information (MI) conveyed by each voxel about each odor (see below). As low MI values (i.e., minimal mutual dependence between the distribution of responses and odor) reflect indiscriminant or random responses to all odors, voxels with bottom 10% MI values in a given ROI were excluded ^42^. Voxel-based tuning was defined by the odor mixture eliciting the largest beta (i.e., optimal odor). As such, each of the remaining voxels was classified into one of five odor classes. In line with animal tuning analysis ^12, 66^, we examined the voxels tuned to the neighboring odors (nCSs and nCSt) of the CS before conditioning and measured their tuning shift to the CS (relative to the neighboring nCS odor/nCSm) after conditioning. Accordingly, we derived a tuning shift index (TSI) for Day 1 and Day 9 post-conditioning: TSI = (% CSs – % nCSm) + (% CSt – % nCSm), reflecting the % of initially nCSs/nCSt voxels that became tuned to the neighboring CSs/CSt, respectively, relative to the % of initially nCSs/nCSt voxels that became tuned to nCSm.

##### MI calculation

First, we converted the beta values into a discrete variable (*B*) by dividing the range of betas into a set of equidistant bins (*b*). The size of the bins was determined by Freedman-Diaconis’ rule [bin size = (max(*B*) – min(*B*))/2*IQR*n^-1/3^], where n is the number of trials (*n* = 75). We selected the median bin size of all voxels within an ROI based on the pre-conditioning data and held it constant for the post-conditioning sessions (Day 1 and Day 9). Next, we computed for each voxel the entropy of (discretized) responses (*B*) as follows:

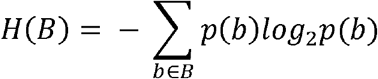

where *p*(*b*) is the proportion of trials whose responses fall into bin *b*. Then, we computed conditional entropy *H*(*B*|*o*), the entropy of responses given knowledge of the odor condition, as follows:

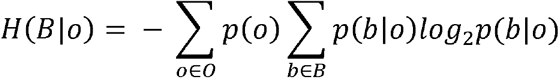

where *p*(*b*|*o*) is the proportion of trials falling into bin *b* when responding to a certain odor (*o*). The index of *MI*(*B*; *O*), i.e., the amount of information a voxel conveys for an odor, was calculated as the reduction in entropy of responses given knowledge of the odor condition:

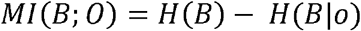

### Statistical analysis

Using analyses of variance (ANOVAs) of Odor (five mixtures) and Time (Day 1 Post and Day 9 Post), we performed trend analysis over the odor continuum on risk ratings and ODT response to capture the warping of affective and perceptual spaces by conditioning. We hypothesized that affective learning via conditioning would change the baseline neutral trend to an ascending safety-to-threat trend (Fig. 2a). We further hypothesized that differential conditioning would enhance perceptual discrimination of the CS, expanding odor quality distances between the CS and their neighboring odors; resulting changes in odor quality space (i.e., differential CS endorsement rates; Post - Pre) would conform to a cubic trend (Fig. 2b). As for the neural mechanisms, i.e., enhanced pattern separation between the CS and similar (neighboring) nCS and tuning shift towards the CS, we conducted ANOVAs of ROI (APC/PPC) and Time (Day 1 Post and Day 9 Post) on differential PSI scores and TSI scores, respectively. Finally, we examined modulatory effects of anxiety using Pearson’s correlation of BIS scores with behavioral and neural effects of conditioning. Significance threshold was set at *P* < 0.05. Given the clear *a priori* hypotheses, one-tailed tests were accepted and are explicitly noted in the Results (two-tailed tests are not explicitly noted). To protect for Type I error, only significant effects in the ANOVAs were followed up with hypothesis testing. Correlational analysis with anxiety involved multiple tests, which were corrected using the false discovery rate (FDR) criterion (i.e., FDR *P* < 0.05).

**Fig. 2.**
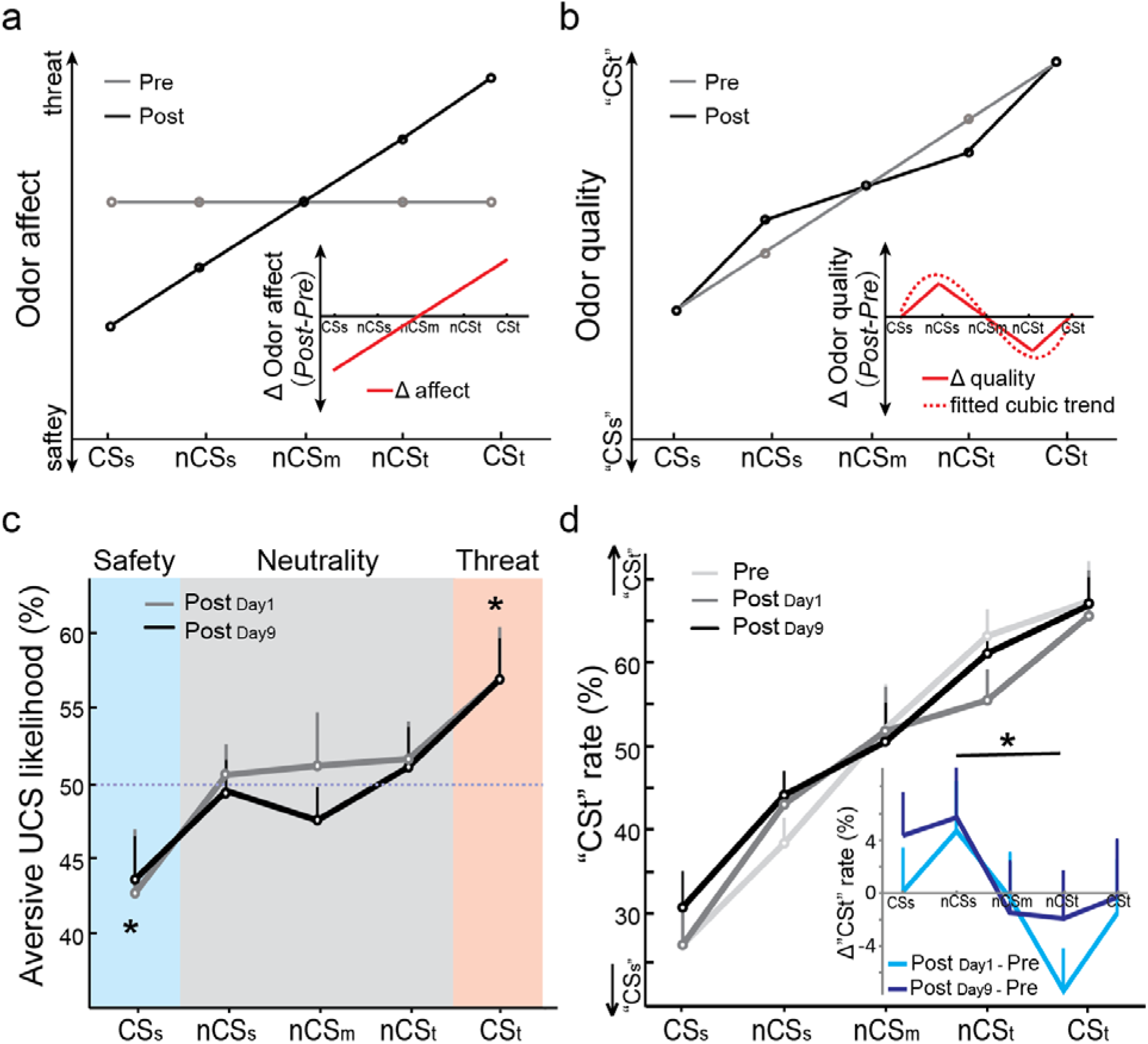
Behavioral effects of olfactory conditioning. (a) Hypothetical affective space over the odor continuum: the initial neutral baseline (gray line) would change to an ascending safety-to-threat line (black line) after acquiring affect (safety/threat) through differential conditioning. Inset shows changes in odor affect over the continuum via conditioning, which conforms to a linear trend. (b) Hypothetical perceptual (quality) space over the odor continuum: the initial ascending trend (gray line; tracking the linear increase in the proportion of CSt) would be warped after conditioning due to expanded distances (i.e., increased perceived dissimilarity) between the CS (CSt/CSs) and neighboring nCS (nCSt/nCSs) (black line). Inset illustrates such changes (Post – Pre) in perceived odor quality (solid line), which can be fitted by a cubic trend (dotted line). (c) Empirical risk ratings (likelihood of aversive UCS) on both days conformed to the predicted profile of differential conditioning: below-chance risk for CSs and above-chance risk for CSt. Risks for the three intermediate mixtures remained chance-level (50%; indicated by the dotted line). (d) Empirical 2-AFC ODT performance (“CSt” responses rate) over the CSs-to-CSt continuum conformed to a linear trend before conditioning, which was warped after conditioning. Inset illustrates differential “CSt” rates (Post - Pre) over the odor continuum on Day 1 and Day 9, which largely conformed to the hypothesized trend, with the nCSt odor less endorsed as “CSt” and the nCSs odor more endorsed as “CSt” (i.e., less as “CSs”). Error bars represent s.e.e. (individually adjusted s.e.m.). *: *P* < .05.

## RESULTS

### Behavioral effects

#### Affective appraisal

Odor valence ratings (on a VAS of 0-100) acquired at the pre-experiment screening indicated neutral affective values for the five odors, conforming to a flat neutral baseline over the odor continuum [*P* = 0.416; Mean (SD) = 50.6 (19.7)]. Risk ratings of the odors (i.e., the likelihood of UCS following a given odor) were acquired post-conditioning on Day 1 and Day 9, which demonstrated a strong ascending linear trend over the odor continuum (*F*_1,30_ = 6.99, *P* = 0.013; Fig. 2c). There was no Odor-by-Time interaction (*P* = 0.908), suggesting equivalent trends for immediate and delayed ratings. Indicating acquired threat and safety value, CSt and CSs had maximal and minimal risk ratings, respectively (CSt vs. CSs: *t*_31_ = 3.02, *P* = 0.005), deviating from the neutral level (50%) in opposite directions (CSt: *t*_31_ = 2.67, *P* = 0.012; CSs: *t*_31_ = −2.40, *P* = 0.023). Ratings for the three nCS odors remained neutral on both days (49.3 - 51.3%; all *P* values > 0.581) and comparable to each other (*F*_1,31_ = 0.14, *P* = 0.708). Nonetheless, they differed from ratings for CSt and CSs in opposing directions (nCSt vs. CSt: *t*_31_ = −2.41, *P* = 0.022; nCSs vs. CSs: *t*_31_ = 1.92, *P* = 0.032, one-tailed). Finally, we correlated anxiety with the changes in risk ratings and observed no significant correlation (all *P* values > 0.252). Together, while inducing limited generalization to the nCS, differential conditioning successfully produced a threat and a safety CS, respectively, which persisted till Day 9.

#### Perceptual discrimination

Consistent with the linear odor morphing continuum, baseline ODT performance conformed to a strong linear trend of increasing endorsement of the dominant odor of the CSt (i.e., “CSt” rate), *F*_1,31_= 79.62, *P* < 0.0001 (Fig. 2d). Consistent with our hypothesis (Fig. 2b), an ANOVA of Odor (five odors) and Time (Day 1/Day 9) on differential “CSt” rates (Post – Pre) showed a cubic trend (*F*_1,31_ = 3.16, *P* = 0.043 one-tailed). Like risk ratings, there was no Odor-by-Time interaction (*P* = 0.405), suggesting equivalent changes for Day 1 and Day 9. To test that this cubic trend was driven by perceptual discrimination between CS and neighboring nCS, we then performed a follow-up ANOVA of Odor (nCSt/nCSs) and Time (Day 1/Day 9) on differential “CSt” rate. We observed an effect of Odor (*F*_1,31_ = 5.19, *P* = 0.030), confirming that the “CSt” endorsement rate decreased for nCSt (i.e., less similarity/greater discrimination with CSt) and increased for nCSs (i.e., less similarity/great discrimination with CSs) from pre-to post-conditioning. Again, this ANOVA showed no Odor-by-Time interaction (*P* = 0.427). Correlation analysis between anxiety (BIS scores) and these difference scores showed no significant correlation (all *P* values > 0.264). In sum, these results indicate that differential conditioning warped odor quality space and particularly, expanded distances (i.e., enhanced perceptual discrimination) between CS and neighboring nCS.

### Neural effects

#### Pattern separation

ANOVAs of ROI (APC/PPC) and Time (Day 9/Day 1) on differential PSI (Post – Pre) showed a main effect of Time, *F*_1,30_ = 4.30, *P* = 0.047, but no effect of ROI (*P* = 0.296) or interaction (*P* = 0.271). This Time effect was due to significant PSI increase in the olfactory (APC and PPC) cortex (*t*_30_ = 1.914, *P* = .033 one-tailed) on Day 1, in contrast to no PSI increase on Day 9 (all *P* values > 0.366; Fig. 3a & b). Nonetheless, correlation analysis with BIS scores indicated a significant correlation between anxiety and olfactory cortical PSI increase on Day 9 (*r* = 0.401, *P* = 0.025, FDR *P* < 0.05; Fig. 3c), indicating lasting olfactory cortical pattern separation among anxious individuals. Notably, as illustrated in Fig. 3, the significant (simple or correlational) effects above were comparable for the APC and PPC (all *P* values < 0.05 one-tailed).

**Fig. 3.**
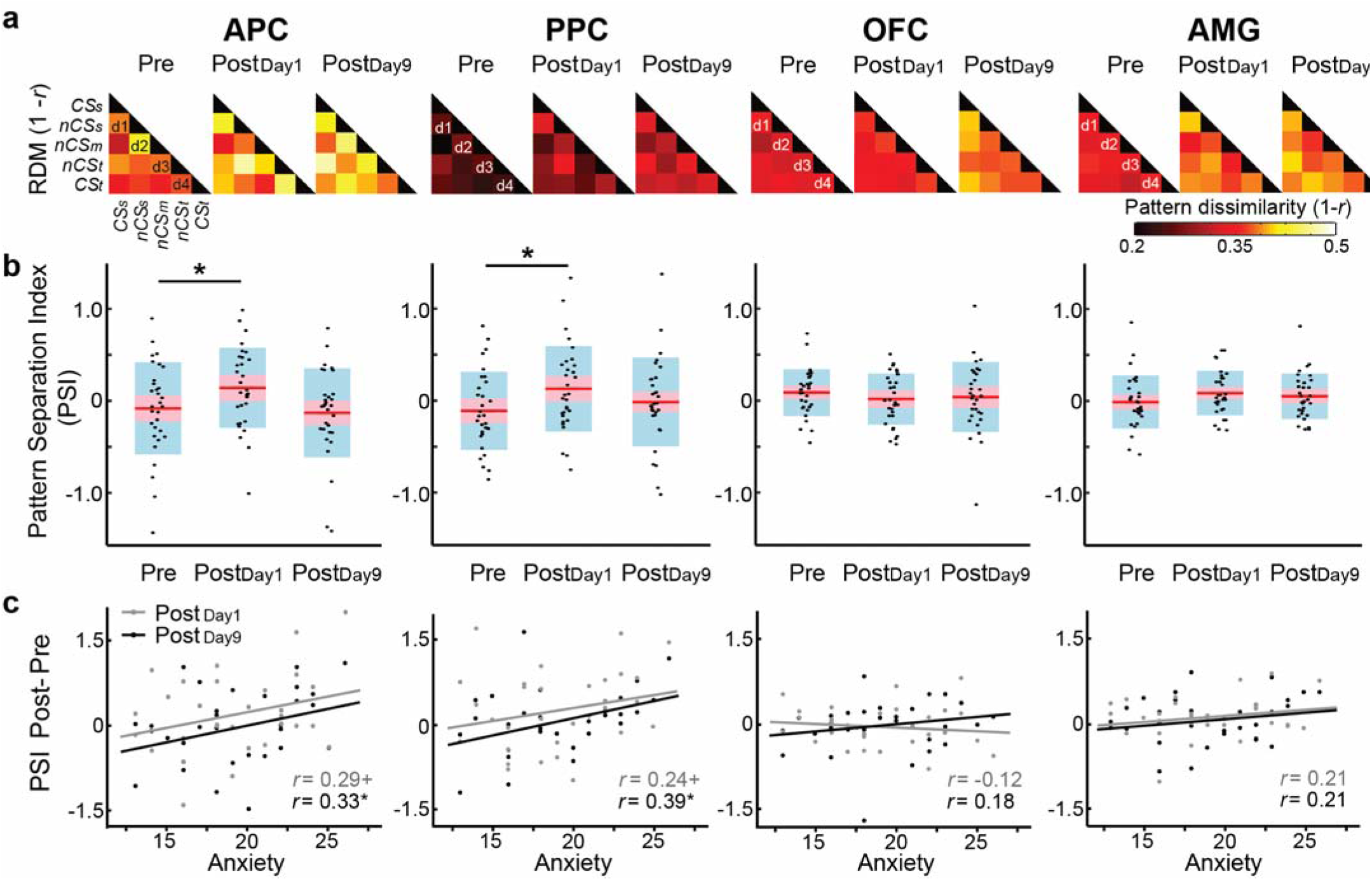
Olfactory cortical pattern separation between CS and neighboring nCS odors. (a) Group-average representational dissimilarity matrices (RDMs) for APC, PPC, OFColf and AMG at each phase. Each cell of the matrix indexes pattern dissimilarity (1-*r*), reflecting pattern separation, for a given odor pair. Cells right off the diagonal index pattern separation between neighboring odors: CSs and nCSs (d1), nCSs and nCSm (d2), nCSm and nCSt (d3), and nCSt and CSt (d4). Based on that, we derived a Pattern Separation Index (PSI) for the CS and the neighboring nCS [PSI = d1 + d4 – (d2 + d3)]. (b) PSI for each ROI at pre-, Day 1, and Day 9 post-conditioning. Both APC and PPC demonstrated increased PSI from pre- to post-conditioning on Day 1, but not on Day 9. Center red line = group mean; red and blue boxes = 95% confidence interval and mean ± 1 SD, respectively. (c) Correlations between conditioning-induced PSI changes and anxiety. PSI changes on Day 9 (vs. Pre) in the APC and PPC correlated positively with anxiety. *: *P* < 0.05; +: *P* < 0.1.

#### Tuning shift

We then examined tuning shift towards CS in the olfactory cortex, i.e., whether voxels initially tuned to neighboring odors of the CS (i.e., nCSt and nCSs) became maximally responsive to the CS after conditioning ^12, 66^. At baseline, there was no difference in % of voxels tuned to the five odors in any of the four ROIs (all *F* values < 1.88, *P* values > 0.125), suggesting evenly distributed tuning for the five odors across the morphing continuum. An ANOVA of ROI (APC/PPC) and Time (Day 9/Day 1) on tuning shift towards CS (TSI scores) revealed a significant ROI-by-Time interaction (*F*_1,30_ = 6.99, *P* = 0.013) and no main effect of ROI (*P* = 0.379) or Time (*P* = 0.612). Specifically, the interaction effect was driven by significant TSI in PPC on Day 9 (*t*_30_ = 3.00, *P* = 0.005; FDR *P* < 0.05) but not on Day 1 (*P* = 0.802) or in the APC on either day (all *P* values > 0.269; Fig. 4a & b). Importantly, correlational analysis with BIS scores again indicated a strong positive correlation between anxiety and TSI, albeit only for Day 9 TSI in the PPC (*r* = 0.44, *P* = 0.014; FDR *P* = 0.056; for the APC and Day 1 PPC TSI, all *P* values were > 0.210), suggesting that this delayed PPC tuning shift was particularly prominent in anxious individuals (Fig. 4c).

**Fig. 4.**
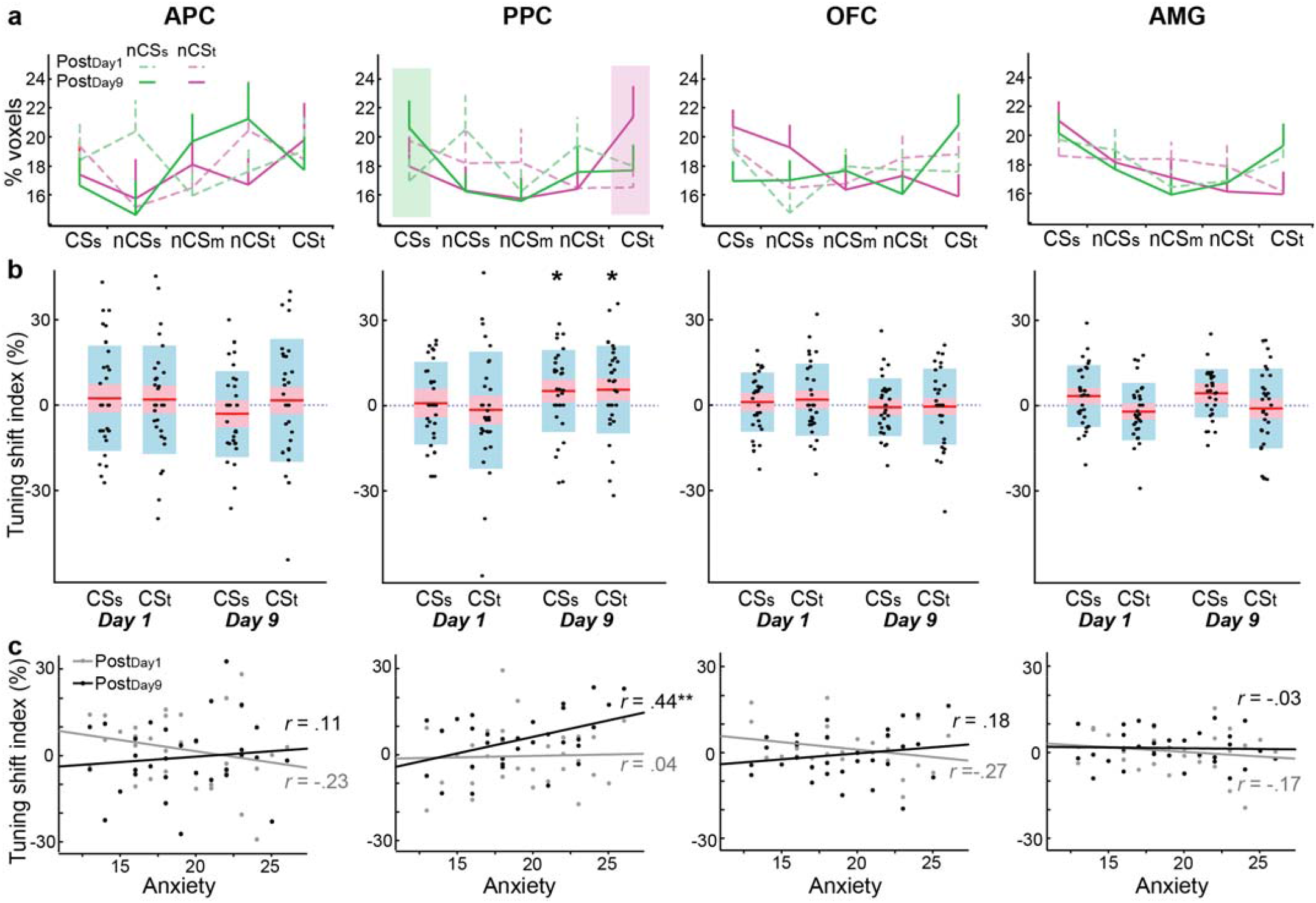
Olfactory cortical tuning shift towards the CS. (a) Day 1 (dashed lines) and Day 9 (solid lines) post-conditioning tuning profiles of nCSs (green) and nCSt (pink) voxels (respectively tuned to nCSs and nCSt at the baseline). In PPC on Day 9, the nCS voxels exhibited a strong tuning preference for their respective CS: highest % of nCSs voxels tuned to CSs (shaded in green) and highest % of nCSt voxels tuned CSt (shaded in pink). (b) Tuning shift index (TSI; % of nCS voxels towards respective CS vs. the middle nCS) on Day 1 and Day 9 post-conditioning. On Day 9, PPC showed significant TSI for both nCSs and nCSt voxels towards their respective CS (CSs and CSt, respectively). The dotted line indicates zero tuning shift (TSI = 0). Center red line = group mean; red and blue boxes = 95% confidence interval and mean ± 1 SD, respectively. (c) Correlations between anxiety and tuning shift towards CS (collapsed across nCSs and nCSt). Day 9 TSI in the PPC correlated positively with anxiety. \**P* < 0.05; \*\**P* < 0.01.

#### Supplemental analysis

We then explored pattern separation and tuning shift in the amygdala and olfactory OFC. As for pattern separation, the amygdala showed a marginal increase in PSI on Day 1 (*t*_30_ = 1.48, *P* = 0.075 one-tailed) but not on Day 9 (*P* = 0.413; Fig. 3a & b). The PSI scores on neither day were correlated with anxiety (all *P* values > 0.252; Fig. 3c). The olfactory OFC showed no PSI increase nor correlations of PSI increase with anxiety (all *P* values > 0.292). As for tuning shift, TSI in the amygdala and olfactory OFC showed no significant tuning shift on either day (All *P* values > 0.169) nor correlation with anxiety (All *P* values > 0.144; Fig. 4). Therefore, in contrast to the primary olfactory cortex, these two regions failed to exhibit clear pattern separation or tuning shift following conditioning.

## DISCUSSION

Growing evidence from aversive conditioning, especially in animals, has compelled the expansion of the “fear circuit” to incorporate the sensory cortex as a key site for fear memory. Here, among human subjects, we demonstrated affective and perceptual learning and memory via conditioning. More importantly, we revealed immediate and lasting pattern separation and late-onset yet lasting tuning shift in the human olfactory cortex, particularly in anxious individuals. These findings provide first mechanistic insights into long-term associative plasticity in the human sensory cortex, highlighting an evolutionarily conserved sensory cortical system of fear memory that could underpin anxiety pathology.

Differential conditioning is known to promote divergent conditioned responses to the (threat and safety) CS, while minimizing conditioning generalization and facilitating CS discrimination (especially from similar stimuli) ^22, 41, 67, 68^. Using affective appraisal (i.e., risk ratings), we established differential affective learning and memory for CSt and CSs and minimal generalization to the nCS (i.e., intermediate odors in the odor morphing continuum). Using an odor discrimination task, we also demonstrated enhancement in perceptual discrimination between the CS and neighboring nCS. Notably, by parametrically morphing odor mixtures to map a linear olfactory continuum, we were able to demonstrate that such affective and perceptual learning/memory resulted in the reorganization of affective and perceptual spaces. Specifically, both affective and perceptual distances were expanded between the CS and their neighboring nCS and compressed between the nCS and their nCS neighbor (i.e., the middle nCS/nCSm). The paralleled affective and perceptual reorganization echoes the idea that acquisition and generalization of fear response tracks the perceptual distance between the CS and nCS ^69, 70^, and perceptual analysis plays an important role in conditioning generalization and specification ^46^.

Paralleling these affective and perceptual expansions (between the CS and neighboring nCS) was enhancement in pattern separation (between the CS and neighboring nCS) in the primary olfactory cortex (both APC and PPC). Importantly, in support of its role in long-term fear memory, this enhanced pattern separation persisted till Day 9, especially in anxious subjects who are known to have heightened fear/threat processing. The human PPC is considered a critical site for storing olfactory sensory representation to support basic odor object encoding ^71, 72^. Our previous study with fMRI recordings immediately after conditioning has revealed immediate enhancement in PPC pattern separation between the CS and its similar nCS ^22^. Here, the lasting enhancement in pattern separation in PPC highlights the enduring differentiation of olfactory representation of the CS (vs. similar nCS), potentially underpinning the long-term memory of acquired threat/safety value. The comparable APC pattern separation here replicates animal findings of decorrelated APC responses to the CS and similar nCS after conditioning ^28, 41^. The human APC is thought to support olfactory attention and arousal ^72^. We surmise that in line with affective learning and memory, this APC pattern separation could underpin the acquired affective value and consequent arousal and attentional response to the CS (vs. similar nCS).

Conditioning also generated tuning shift to the CS, which emerged in the PPC (but not in the APC), highlighting its association with odor object encoding. Interestingly, this PPC tuning shift did not occur immediately and was observed on Day 9 only. This temporal pattern is largely consistent with animal evidence: sensory cortical tuning shift is relatively weak in magnitude and specificity immediately after conditioning but progresses in specificity and strengthens over time (days and weeks) ^14^. Also consistent with this temporal profile, a previous study from our lab showed that enhanced response to the CS in the primary visual cortex (V1/V2; reflective of enhanced tuning of the CS) emerged at the retention (15-day) test but not immediately after conditioning ^27^. As aforementioned, sensory cortical tuning shift to the CS represents associative representational plasticity, by which the sensory cortex updates and stores the new representation/encoding of the CS ^6, 14^. Along this line, tuning shift in the human PPC here indicates the development of long-term memory in the sensory cortex to underlie the “acquired associative representation” of the CS ^6, 14^.

In comparison, the amygdala and OFC exhibited no clear evidence of enhanced pattern separation or tuning shift by conditioning. It is important to note that we analyzed pattern separation/tuning shift expressly along a physical dimension (i.e., odor-morphing continuum) to elucidate changes in neural representation of sensory input. Therefore, the null effects in the amygdala and OFC would not rule out associative plasticity in other, abstract dimensions (e.g., valence or value). In fact, previous research comparing (immediate, appetitive) conditioning effects in the rodent piriform cortex and OFC has revealed sensory-based plasticity in the former and value/rule-based plasticity in the latter (e.g., ^73^). Similarly, in humans, previous research of (both appetitive and aversive) conditioning has underscored value-based (vs. sensory-based) pattern separation in the OFC and amygdala ^22, 74, 75^.

Finally, we demonstrated modulatory effects of anxiety in these sensory cortical mechanisms. That the anxiety effects were most prominent on Day 9 emphasizes its pivotal role in long-term fear memory. Specifically, while pattern separation appeared immediately, it was present in anxious individuals only on Day 9, suggesting that anxiety helps to resist memory decay. As for tuning shift that emerged on Day 9 only, anxiety amplified its strength, suggesting that anxiety facilitates the development of fear memory. This modulatory effect of anxiety echoes our previous demonstration of a positive association between anxiety and long-term associative plasticity in the visual cortex^27^. The fear conditioning model of anxiety has strong conceptual justification and compelling animal evidence ^2, 3^, but human laboratory evidence has indicated only modest modulatory effects of anxiety ^76^. Tapping into sensory cortical plasticity, the current study was able to demonstrate the importance of anxiety in aversive conditioning. Moreover, as such associative sensory cortical plasticity figures importantly in long-term fear memory, the current finding lends credence to the theory of a hyperactive sensory-bound representation system of threat memory (“S-memory”) in anxiety ^77, 78^. This hyperactive sensory memory system of fear can account for hallmark symptoms of intrusive memories in posttraumatic stress disorder (PTSD), which are laden with vivid sensory fragments of trauma and can be readily triggered by simple sensory cues ^79, 80^. In keeping with that, a clinical study from our group showed that sensory cortical disinhibition/overactivation could mediate excessive olfactory trauma memory in PTSD^81^. Taken together, the anxiety effects here add to the growing evidence in the literature, advocating for a sensory mechanism—exaggerated sensory cortical representation of threat—in the pathogenic model of anxiety ^82^.

To conclude, the current study advances the burgeoning human literature of sensory cortical plasticity via aversive conditioning, promoting a multi-system conceptualization of fear ^83^ and an expanded fear circuit in humans ^9^. By identifying pattern separation and tuning shift in the human sensory cortex, the current study specifies CS representation in the sensory cortex as a key component of long-term fear memory. Importantly, that this process is heightened in anxiety sheds new light on the neuropathophysiology of anxiety and related disorders and identifies a new target for clinical intervention.

## DATA AVAILABILITY

Anonymized data and code created for the study will be available in a repository (NIMH Data Archive) upon publication. Data Type: anonymized T1 images, EPI scans, behavioral data, respiratory recordings, and data analysis scripts.

## CODE AVAILABILITY

The data analysis scripts will be available in a repository (NIMH Data Archive) upon publication.

## AUTHOR CONTRIBUTIONS

Y.Y., L.N., and W.L. designed the experiment; Y.Y., L.N., K.C. collected data; Y.Y., L.N., and W.L. performed analysis; W.L., Y.Y., L.N., and K.C. wrote the manuscript.

## SUPPLEMENTAL MATERIAL

### Supplemental Results

#### Baseline odor ratings

We performed analyses on odor ratings to exclude confounds related to inherent odor stimulus differences. Baseline ratings for all five odor mixtures on valence, intensity, familiarity, and pungency were submitted to separate repeated-measures ANOVAs, which revealed no significant difference among five odor mixtures on any of the scales (all *F* values < 1.61, all *P* values > 0.182).

#### Respiration

We also examined respiration parameters during the 2-AFC ODT, including peak amplitude, peak latency, and sniff inspiratory volume. ANOVAs (Odor X Time) on these sniff parameters revealed no effects of odor or odor-by-time interactions (all *P* values > 0.095). These results thus ruled out variations in sniffing as potential confounds.

